# Interpretable improving prediction performance of general protein language model by domain-adaptive pretraining on DNA-binding protein

**DOI:** 10.1101/2024.08.11.607410

**Authors:** Wenwu Zeng, Yutao Dou, Liangrui Pan, Liwen Xu, Shaoliang Peng

## Abstract

DNA-protein interactions exert the fundamental structure of many pivotal biological processes, such as DNA replication, transcription, and gene regulation. However, accurate and efficient computational methods for identifying these interactions are still lacking. In this study, we propose a novel method ESM-DBP through refining the DNA-binding protein (DBP) sequence repertory and domain-adaptive pretraining based the protein language model (PLM). Our method considers the lack of exploration of general PLM for DBP domain-specific knowledge, so we screened out 170,264 DBPs from the UniProtKB database to construct the model that more suitable for learning crucial characteristics of DBP. The evaluation of ESM-DBP is systematically performed in four different DBP-related downstream prediction tasks, i.e., DNA-binding protein, DNA-binding residue, transcription factor, and DNA-binding Cys2His2 zinc-finger predictions. Experimental results show that ESM-DBP provides a better feature representation of DBP compared to the original PLM, resulting in improved prediction performance and outperforming other state-of-the-art prediction methods. In addition, ESM-DBP incorporates the integrated gradient algorithm for interpretable analysis, which usually ignored in the previous methods. It reveals that ESM-DBP possesses high sensitivity to the key decisive DNA-binding domains. Moreover, we find that ESM-DBP can still perform well even for those DBPs with only a few similar homologous sequences, and this generalization performs better than the original PLM. The data and standalone program of ESM-DBP are freely accessible at https://github.com/pengsl-lab/ESM-DBP.

## INTRODUCTION

Proteins and DNA interact throughout most of life activities. Most typically, transcription factors (TFs) direct gene expression by orchestrating the transcription process [1]. In addition, DBPs like DNA polymerase, DNA topoisomerase, and histone are also involved in a variety of biological functions to form a complex system. These DBPs often serve as pivotal contributors to disease etiology, e.g., hotspot mutations in the transcription factor IKZF3 lead to B cell neoplasia [2]; deletion of the repressor element 1-silencing transcription factor is associated to neurological disease such as Alzheimer’s disease [3]. Exploring the intrinsic mechanisms of protein-DNA interaction contributes to the understanding of disease pathogenesis. Biological experiment-based approaches such as systematic evolution of ligands by exponential enrichment (SELEX), chromatin immunoprecipitation (ChIP), X-ray crystallography, and nuclear magnetic resonance (NMR) have made outstanding contributions to the study of DNA-protein interactions over the past decades. However, their utility is inherently constrained by high costs and time-intensive protocols, necessitating the incorporation of computational approaches to address this challenge.

In the last decade, deep learning-based techniques have been widely used in studies related to protein-DNA interaction prediction [4-6]. Nevertheless, most of such methods rely heavily on limited primary sequence in the training datasets and high-quality multiple sequence alignment (MSA) information, thereby restricting their generalizability and accuracy. The emergence of large-scale language models (LLMs) provides a compelling opportunity to overcome these limitations. In the field of natural language processing (NLP), the well-trained LLMs adeptly capture intricate word dependencies and contextual nuances, thus enabling it to communicate like a human in a dialogue. The same reasoning can be naturally extended to protein sequence, structure, and function studies. Generally speaking, the tertiary structure of a protein is encoded in intricate sequence space by 20 standard amino acids, and then fundamentally dictates most of the biochemical functions. The PLM based on vast primary sequences can capture the complex mapping relationships between sequence, structure, and function. Thanks to the rapid development of sequencing technology in the post-genome era, public databases such as UniProt [7] store huge amounts of protein sequences belonging to various species, which are the basis for building large-scale protein language model (PLM). Using PLM to explore the relationship between protein sequence, structure, and function has received focused attention from many bioinformatics researchers [8-13]. For example, by performing unsupervised pretraining on 250 million primary sequences, Rives *et al*. constructed a PLM with 650 million parameters named ESM-1b which contains knowledge about biological properties in its feature representation space, and performs accurate protein secondary structure, residue contact map, and mutation effect predictions [14]. In recent studies [15-18], high-precision protein 3D structure can be inferred using PLM, especially on orphan proteins where the performance exceeds that of AlphaFold2 [19] and RoseTTAFold [20]. Taking ESM2 as an example [15], the PLM with 15 billion parameters constructed using ∼65 million non-redundant sequences sampled from ∼43 million sequences in the UniRef50 database [21] for pretraining performs the high 3D structure prediction accuracy at the atomic-level. Due to the highly conserved and specific nature of DBP functional sequences, it is feasible to use PLM for protein-DNA interaction-related task prediction. For instance, Liu *et al*. developed a DNA-binding residue (DBS) prediction method called CLAPE-DB [22] by fine-tuning the PLM ProtTrans [23], and shows well performance. In ULDNA [24], Zhu *et al*. leveraged an LSTM-attention neural network which embedded with sequence representations of three unsupervised PLMs, i.e., ESM2, ProtTrans, and ESM-MSA [25], to deduce the DNA-binding sites. Roche *et al*. [26] proposed an equivariant deep graph neural networks (GNN)-based predictor named EquiPNAS which employed multiple structure and sequence feature including embedding from ESM2 for accurate nucleic acid-binding residue prediction. In our previous study ESM-NBR [27], a BiLSTM-based multi-task learning framework with ESM2 sequence embeddings as input features is used to achieve fast and accurate nucleic acid-binding residue prediction. What is certain is that the rise of PLM points in a new direction for the computation-based studies of protein sequence, structure, and function.

Although PLMs have achieved state-of-the-art (SOTA) performance across various tasks pertaining to protein structure and function prediction, as the general language model, it has not paid particular attention to proprietary field knowledge since a wide range of protein functions are hidden in the massive pretraining dataset and no prior knowledge is provided during the pretraining phase. Considering the diversity, sequence specificity, and highly conserved nature of protein functions, the ability of general PLM to characterize sequences with specific functions such as DBP should be under-explored. In addition, owing to the substantial parameters inherent in PLMs, interpretability of the predictive performance is often lacking. Previous study shows that domain-adaptive pretraining can further enhance the performance of LLM within a specific domain by learning the knowledge of the given domain [28]. As carriers of life activities, the multifaceted functions exhibited by proteins are intricately linked to the specificity of their primary amino acid sequences. ESM-1b reports that PLM has the ability to encode the remote homology and the alignment within a protein family. It is therefore plausible to associate PLMs with the capability to further discern latent identification patterns specific to a given function from an extensive repository of annotated protein sequences. In this study, to further exploit the characterization capability of PLM on DBPs, we built a interpretable DBP proprietary PLM based on the universal PLM ESM2 (abbreviated as ESM-DBP) and then evaluated it on four DBP-related downstream tasks, i.e., DBP prediction [29, 30], DNA-binding residue (DBS) prediction [31], transcription factor (TF) prediction [32], and DNA-binding Cys2His2 zinc-finger (DBZF) prediction [33]. To be specific, we first downloaded all ∼4 million sequences annotated DBP up to August 2023 from the UniProtKB database [7]; then, redundant sequences of high similarity as well as those sequences that share high similarity with test protein sequences are removed using the CD-HIT tool [34] with a cluster threshold of 0.4, remaining 170,264 non-redundant sequence as the pretraining data set named UniDBP40; then, to prevent overfitting and retain the basic biological knowledge learned by original ESM2, the parameters of first 29 transformer blocks of the ESM2 model with 650 million parameter are frozen and only the last 4 transformer blocks are updated; finally, the biological feature representation of well-trained model is extracted and applied to four downstream classification tasks by supervised learning. The well-trained ESM-DBP model not only contains the basic biological information that ESM2 has learned from the vast number of primary universal sequences but also has the knowledge of specific DNA-binding patterns learned from DBP sequences. Experimental results on the benchmark dataset show that after domain-adaptive pretraining on the DBPs training data set, the feature representation of the ESM2 model not only further improves the prediction performance of the relevant tasks, but also far outperforms the existing SOTA methods which is mostly based on evolutionary information like Hidden Markov Model Profile (HMM) [35] and Position-Specific Scoring Matrix (PSSM) [36]. Remarkably, even without the fine-tuning phase of supervised learning, most of the DBP families contained different DNA-binding domains (DBD) can still be clearly distinguished by the unsupervised clustering, meaning that the model still spontaneously perceives DBD-specific sequence knowledge without label guidance. Using an interpretable algorithm of integrated gradients, our investigation reveals that the enhanced DBP sequence characterization capability and predictive efficacy of ESM-DBP stem from a notable emphasis on various specific DBDs such as Homeobox and Fork-head. Furthermore, the commendable predictive outcomes observed for DBPs with limited homologous sequences underscore the robust generalization capability of ESM-DBP. These findings suggest that ESM-DBP effectively harnesses the latent knowledge embedded within DBP sequences, thereby amplifying the characterization and predictive prowess of PLM, and provides a new perspective for protein-DNA interaction and other protein function researches. Notably, the efficiency of ESM-DBP can extend seamlessly to vast repositories of unexplored sequences, obviating the need for laborious multiple sequence comparison. Here, we provide the prediction results for all 183,801 human proteins in TrEMBL at https://huggingface.co/zengwenwu/ESM-DBP, aiming to accelerate the process of DNA-protein interaction studies. The data and standalone program of ESM-DBP are freely accessible at https://github.com/pengsl-lab/ESM-DBP.

## MATERIALS AND METHODS

### Benchmark datasets

Five data sets are used in this study, i.e., the pretraining set for the unsupervised DBP domain-adaptive pretraining and the data sets of four downstream prediction tasks (DBP, DBS, TF, and DBZF predictions) for supervised learning.

To continue training the ESM2 model, we first download 4,159,929 annotated DBPs from the UniProtKB database until 2 August 2023; then, the CD-HIT program with a cluster threshold of 0.4 is employed to remove redundant sequences of high similarity, as well as those sequences share high similarity with test protein sequences; finally, the remaining 170,264 non-redundant DBP sequences (abbreviate as UniDBP40) are used as the pretraining data set. The data sets of four downstream prediction tasks are obtained from previous studies, i.e., TargetDBP+ [37], PredDBR [31], TFPred [38], and DeepZF [33], respectively. Concretely, the data set for DBP prediction contains one training set *UniSwiss-Tr* (including 4,500 DBPs and 4,500 non-DBPs) and one independent test set *UniSwiss-Tst* (including 382 DBPs and 382 non-DBPs); the data set for DBS prediction consists of one training set PDNA-543 (including 9,549 DBSs and 134,995 non-DBSs) and one independent test set PDNA-41 (including 734 DBSs and 14,021 non-DBSs); The training (or test) data set for TF prediction contains 416 TFs and 413 non-TFs (or 106 TFs and 106 non-TFs). The data set for DBZF prediction named C-RC contains 834 DNA-binding and 789 non-binding ZFs.

### Domain-adaptive pretraining based on universal PLM

The universal PLM ESM2 is constructed based on ∼65 million non-redundant protein sequences by leveraging masked pretraining technique, achieving the protein 3D structure prediction with atomic-level accuracy. Similarly, the well-trained ESM2 model can be used for protein function prediction due to the conserved and specific nature of protein function in terms of sequence and structure. However, considering the diversity and complexity of protein functions, when we target a specific function like DBP, the consequence of a large training set is that the model is overloaded with redundant information, resulting in field-specific information not being given extra attention. Figure 1B illustrates that the percentage of DBP in the pretraining data set of ESM2, i.e., UniRef50 is only 0.0069%. Here, we aim to improve the sensitivity and attention of PLM to DBP-related knowledge by continuing to train the ESM2 model on DBP sequences in such a way as to perform better DBP sequence characterization and achieve performance improvement in downstream prediction tasks. As shown in Figure 1A, in the original PLM model space, there are a variety of protein attribute information including structure and various functions. Our goal is to make the model focus more on regions that overlap with the DBP field, thus learning DBP field-specific knowledge and then improving the accuracy of DBP-related tasks. Figure 1C demonstrates that UniDBP40 and UniRef50 have both overlapping parts and individual sequences, indicating that ESM-DBP not only enhances the expression of DBP knowledge already available in ESM2 but also learns important discriminatory knowledge of unseen DBP sequences. The ESM2 model includes a total of six versions with parameter counts of 8 million, 35 million, 150 million, 650 million, 3 billion, and 15 billion. The 3 billion and 15 billion versions of ESM2 spend 30 and 60 days on 512 NVIDIA V100 GPUs, respectively. Constrained by computational resources, most researchers could not afford to train the networks of this enormous magnitude, so the ESM2 model with 650 million parameters is used to continue training here. In addition, at least our previous study about nucleic acid-binding residue prediction shows that the performance of a model with large parameters is not necessarily better than that of the model with relatively small parameters [39]. We follow the standard masked pretraining approach as ESM2, by randomly masking 15% of the residues in the input sequence and then predicting them. The ESM2 consists of 33 layers of stacked transformer encoder blocks. Considering that the training dataset of ESM2 is much larger than the dataset of this study, to prevent overfitting, we froze the parameters of the first 29 layers of the ESM2 and updated the weights of only the last four layers of the transformer encoder block as well as the logistic layer for amino acid classification prediction. The first 29 layers retain the important biological property knowledge learned by the original ESM2 from the massive protein primary sequences, and the last four layers learn DBP field-specific information through domain-adaptive pretraining. In the training phase, to solve the problem of the unequal length of protein sequences, the sequences with amino acid number greater than 512 are cut into sub-sequences, and the sequences with amino acid number less than 512 are filled with the token of “pad>“ to the length of 512. The batch size is set to 100. The loss function and optimizer are consistent with ESM2. The model is trained for approximately 51k steps and took about 3 days on four Tesla V100 GPUs with 16G memory.

**Figure 1.**
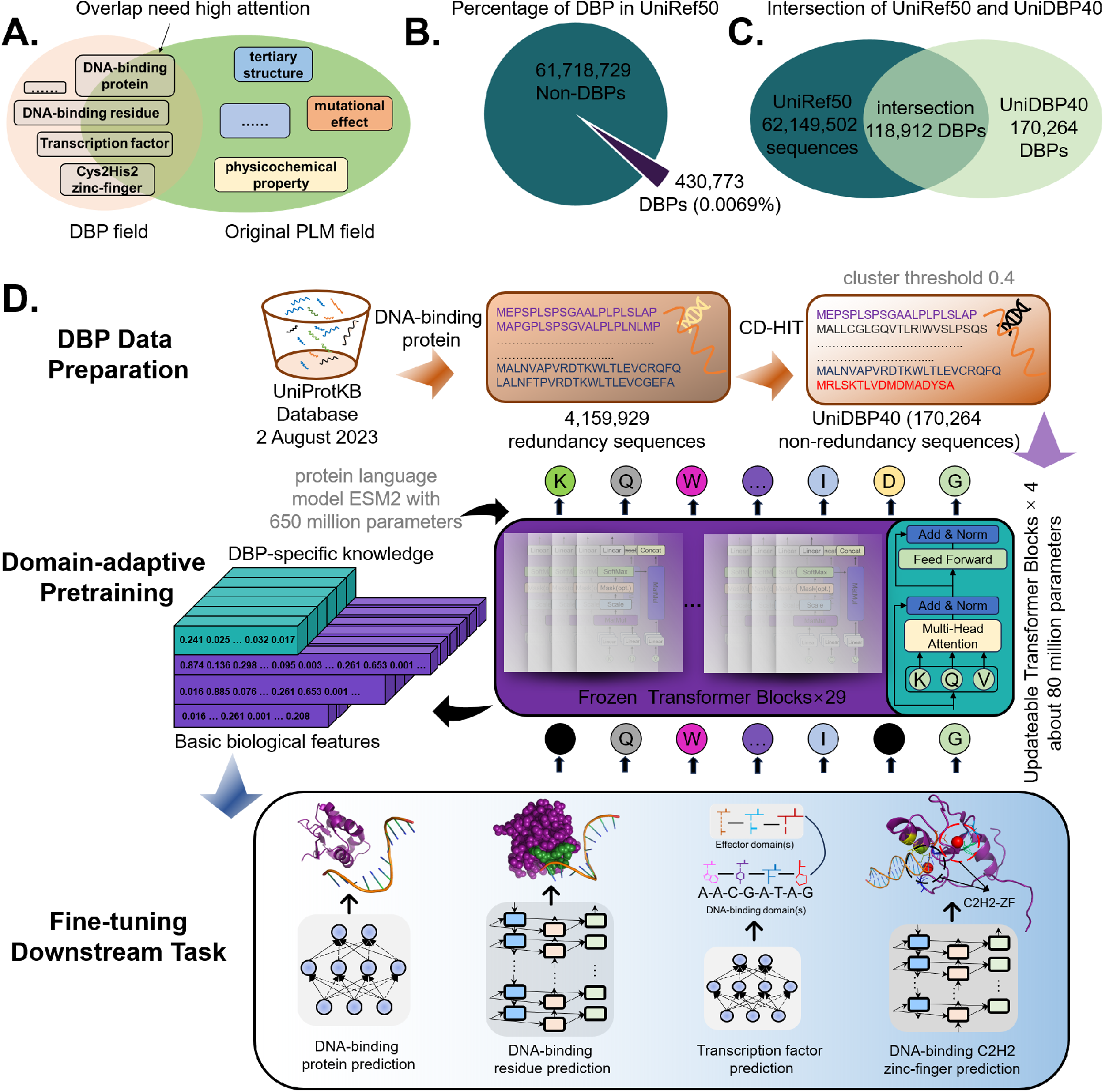
ESM-DBP is a DBP field-specific PLM by domain-adaptive pretraining based general PLM, i.e., ESM2, on the DBP dataset. **(A)**. Data distribution of original ESM2 and DBP domain-specific knowledge; **(B)**. The proportion of DBPs in the original pretraining dataset of ESM2, i.e., UniRef50; **(C)**. Intersection of UniRef50 and the pretraining dataset UniDBP40 in this study; **(D)**. The overall architecture of ESM-DBP.

### Architecture of ESM-DBP

In this study, a DBP field-specific PLM called ESM-DBP based on the ESM2 model is proposed. As shown in Figure 1D, the architecture of ESM-DBP consists of three main steps:

*Step 1. Data Preparation*: a training data set containing 170,264 non-redundant DNA-binding protein sequences is constructed from all DNA-binding sequences in the UniProtKB database until 2 August 2023 using the CD-HIT tool with a cluster threshold of 0.4;

*Step 2. Domain-adaptive Pretraining*: the original ESM2 model with 650 million parameters is pretrained on the DBP training data set and only the parameters of the last four transformer blocks and the logistic layer for classification are updated;

*Step 3. Fine-tuning Downstream Task*: the biological feature representation contained important DBP knowledge of the well-trained model is applied to four downstream DBP-related prediction tasks, i.e., DBP, DBS, TF, and DBZF predictions using the lightweight bidirectional long short-term memory (BiLSTM) and multi-layer perceptron (MLP) networks.

## RESULTS AND DISCUSSIONS

### Role of Domain-adaptive Pretraining on Performance

To observe the impact of domain-adaptive pretraining on the prediction performance, we compare the prediction results of the original ESM2 model and the ESM-DBP for four downstream tasks. In particular, the feature representations of the above two models of size *L* × 1280 are first extracted as input of neural network of four prediction tasks, where *L* denotes the number of amino acids in the protein sequence; then, the prediction results of DBS, DBP, and TF are obtained by independent validation, and the prediction result of DBZF are obtained by 10-fold cross-validation since the DBZF dataset is not divided into training and test sets.

Table 1 shows 7 detailed evaluation indexes of two models. ESM-DBP achieves an overwhelming advantage regardless of the prediction task. Take DBP prediction as an example, the Sen, Spe, Acc, Pre, F_1_, MCC, and AUC values of ESM-DBP are 92.12, 93.43, 93.35, 92.78, 0.927, 0.858, and 0.964, which are 2.62, 3.49, 3.44, 3.06, 3.00, 7.11, and 1.15% higher than those of ESM2, respectively. Moreover, the MCC values of ESM-DBP on DBS, TF, and DBZF predictions are 0.430, 0.924, and 0.349, which are 11.68, 4.17, and 46.02% higher than those of the original ESM2, respectively. Figure 2A shows the ROC curves of ESM-DBP and ESM2 on four prediction tasks. The ESM-DBP has higher AUC values than the ESM2 for all four tasks, especially for DBS prediction where the former achieved an improvement of 5.01% over the latter. In Figure 2B, we know that the ESM-DBP has better predictive results than ESM2 on most samples. Take DBP prediction as an example, out of 381 DBPs (or non-DBPs), there are 308 (or 319) cases in which ESM-DBP has a higher (or lower) prediction probability than ESM2. This result means that for about 80% of the DBPs, ESM-DBP has a better confidence level. Meanwhile, to observe the overall distribution of prediction results for ESM-DBP and ESM2, we make the violin plot of predicted probability for positive and negative samples for each of the four tasks (see Figure 2C). The distribution of the positive samples shows that the overall prediction scores of TF and DBP are significantly higher than those of DBS and DBZF predictions. The main reason for this is that the specific discriminative patterns of DBP and TF from primary sequences are more easily captured by neural networks compared to DBS and DBZF predictions. We also note that the lower end of the DBS positive sample has a higher density, suggesting that the model is not able to make a confident judgment on these residues. Looking more closely, we can see that the ESM-DBP is less dense in this part than the ESM2. This suggests that the PLM after domain-adaptive pretraining mines some of the useful hidden patterns of DNA-binding residue from the large number of DBP sequences. By looking at the internal box plots of the two models on the DBP predictions, the box of ESM2 on the positive (or negative) sample is obviously lower (or higher) than that of ESM-DBP. This shows that ESM-DBP has a better discriminatory ability than original ESM2 both on DBP and non-DBP predictions. Figure 2D visualizes the feature representation of ESM2 and ESM-DBP on the data set of TF prediction before supervised learning using the T-SNE algorithm. By comparison, it is easy to see that the ESM-DBP makes a much clearer distinction for most TFs compared to the original ESM2 both on the test set and the training set. Only a small percentage of TFs are confused with Non-TFs, and most cases can be distinguished by the naked eye from the projection of ESM-DBP. On the contrary, the projection of ESM2 looks worse than ESM-DBP. Many positive and negative samples are so entangled that they are difficult to identify with the naked eye, especially on the training data set. This differential result gives us reason to believe that after domain-adaptive pretraining, PLM spontaneously senses specific discriminatory information of transcription factor function in a large number of DNA-binding protein sequences. Thus, the PLM model is still able to make different representations of TF and Non-TF without being guided by labels. As a conclusion, after retraining on DBP domain-specific data, the PLM mines useful knowledge related to DBP, which enhances the ability of the model to characterize DBP and thus significantly improves downstream prediction task performance.

**Table 1.** Performance comparison of original ESM2 and ESM-DBP on four DBP-related prediction tasks.

| prediction task | model | Sen | Spe | Acc | Pre | F<sub>1</sub> | MCC | AUC |
| --- | --- | --- | --- | --- | --- | --- | --- | --- |
| DBP | ESM2 | 89.76 | 90.28 | 90.24 | 90.02 | 0.900 | 0.801 | 0.953 |
|  | ESM-DBP | **92.12** | **93.43** | **93.35** | **92.78** | **0.927** | **0.858** | **0.964** |
| DBS | ESM2 | 40.32 | **97.17** | **94.34** | **42.77** | 0.415 | 0.385 | 0.859 |
|  | ESM-DBP | **66.62** | 92.89 | 91.58 | 32.90 | **0.440** | **0.430** | **0.905** |
| TF | ESM2 | 96.22 | 92.45 | 92.72 | 94.34 | 0.944 | 0.887 | 0.970 |
|  | ESM-DBP | **97.17** | **95.28** | **95.37** | **96.22** | **0.962** | **0.924** | **0.981** |
| DBZF<sup>a</sup> | ESM2 | 66.42 | 57.41 | 62.04 | **64.55** | 0.664 | 0.239 | 0.673 |
|  | ESM-DBP | **80.33** | **53.23** | **67.16** | 64.48 | **0.715** | **0.349** | **0.711** |

a. The prediction results of DBZF are obtained by 10-fold cross-validation.

**Figure 2.**
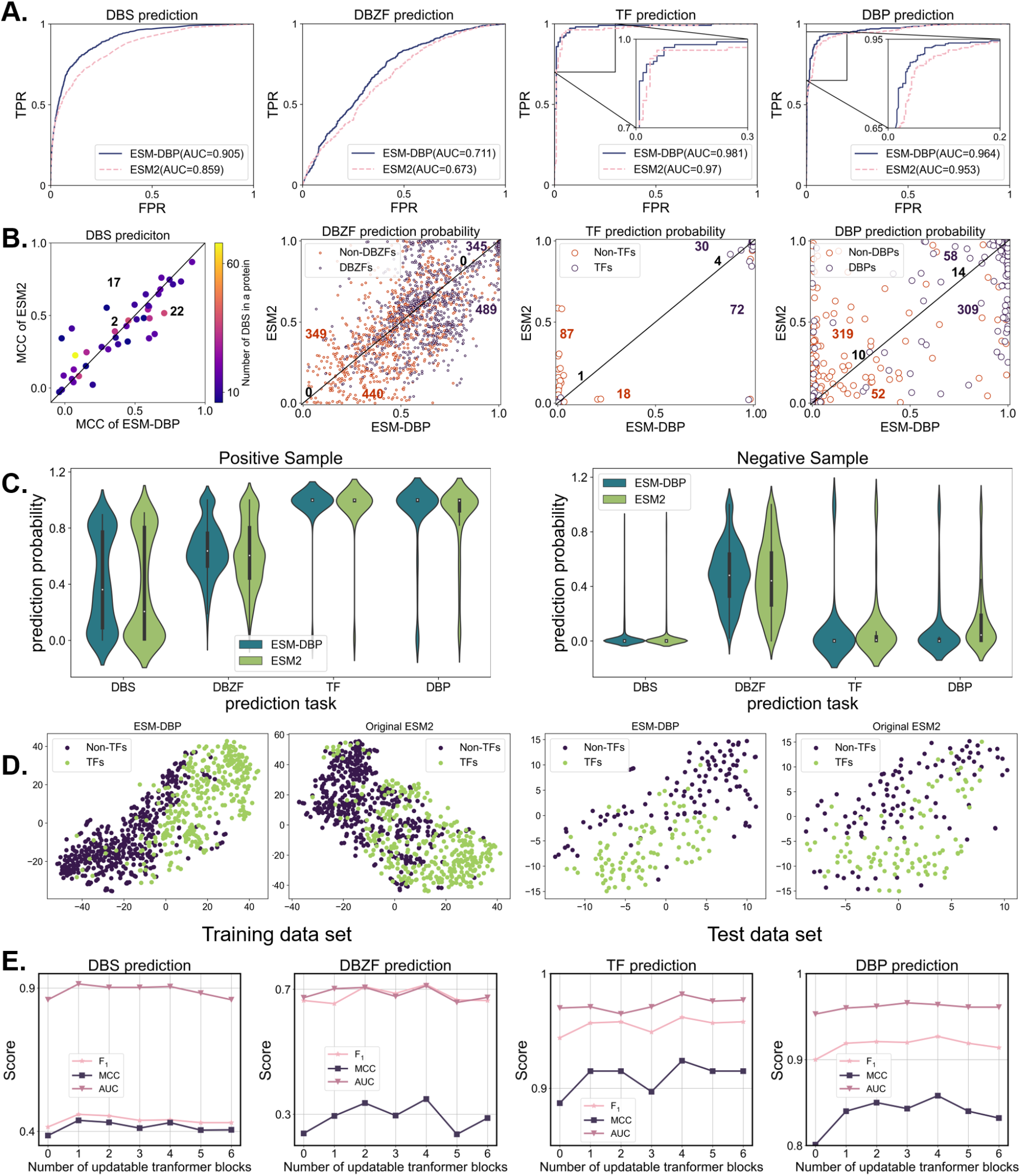
ESM-DBP provides better discrimination than the original ESM2. **(A)**. ROC curves of ESM-DBP and ESM2 on four prediction tasks; **(B)**. Scatterplots of ESM-DBP and ESM2 on four prediction tasks. Each point represents a sample and the number indicates the number of positive or negative samples located in the diagonal, upper, or lower triangle. The axes of the DBS represent the MCC values for each protein and the other three axes indicate the probability of a sample being predicted as positive; **(C)**. Violin plot of the probability of all samples being predicted as positive; **(D)**. T-SNE [40] visualization of feature representations of ESM-DBP and ESM2 on TF data set before supervised learning; **(E)**. Impact of the number of updatable transformer blocks on prediction performance. The number of parameters per layer transformer block of ESM2 is about 20 million. The number of 0 represents the original ESM2.

### Determining the number of parameters to be updated

To achieve highly accurate protein 3D structure prediction, the original ESM2 is constructed on ∼43 million protein sequences, which are almost 150 times the training set of this study. To fit such a large amount of data, the largest ESM2 model has a parameter count of 15 billion. Although the ESM2 model used in this study is a 650M parameter version, given the limited number of sequences in our training set, it is still inappropriate to directly update all parameters in the model. On the one hand, the well-trained PLM contains important knowledge of the underlying biological of proteins learned from a huge number of primary sequences. This critically important biological information will be corrupted leading to deterioration of sequence characterization if all parameters are updated. On the other hand, using such a huge neural network to fit a relatively small amount of training data will undoubtedly lead to overfitting and thus learning redundant noise, which can also lead to worsening prediction performance. In this study, to take advantage of both the underlying biological feature attributes already learned by the original ESM2 and the specificity knowledge in the DBP sequences, we freeze most of the transformer blocks in ESM2, leaving only a small number of updateable parameters. However, it is not an easy task to determine the proper number of updatable parameters to fit the existing training data, especially considering the huge number of parameters of the original ESM2 model and the complex non-linear relationship between protein sequences and DBP functions.

Here, start with the original ESM2, we gradually increase the number of updatable transformer blocks with the step of 1, and determine the final model structure by observing performance on downstream tasks. From Figure 2E, two basic facts can be observed: first, the model outperforms the original ESM2 even when only one layer of transformer block parameters is updated; second, the model achieves a more outstanding performance on all tasks when the number of updatable transformer blocks is 4. Therefore, the number of adjustable transformer blocks for the final ESM-DBP model in this study is set to 4. The number of parameters per layer transformer block of ESM2 is about 20 million, so the number of parameters to be updated is about 80 million.

### ESM-DBP outperforms SOTA methods over four prediction tasks

To further assess the performance of the proposed ESM-DBP in predicting the four downstream tasks, 22 SOTA prediction methods, including 8 for DBP (MsDBP [42], iDNAProt-ES [43], TargetDBP [44], TargetDBP+ [37], TPSO-DBP [29], iDRBP_MMC [45], iDRBP_ECHF [46], and iDRPro-SC [47]), 10 for DBS (NCBRPred [48], CLAPE-DB [22], DRNAPred [49], iDRNA-ITF [50], iProDNA-CapsNet [51], DNABind [52], TargetDNA [53], DNAPred [54], COACH [55], and PredDBR [31]), 3 for TF (TFPred [38], DeepTFactor [32], and Wang *et al*’s work [56]) and 1 for DBZF (DeepZF [33]) are employed as control. Figure 3 shows the overall comparison results.

**Figure 3.**
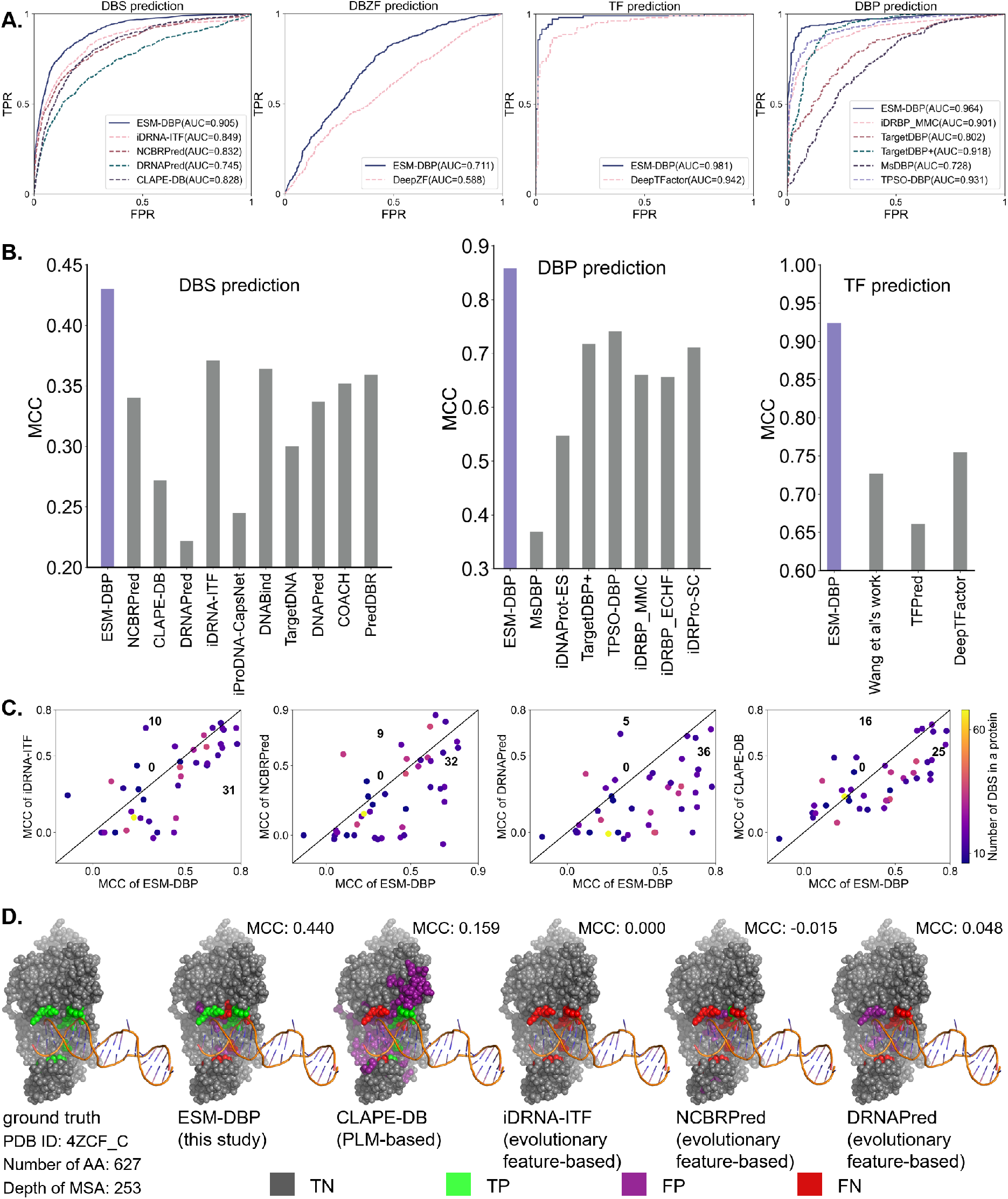
ESM-DBP outperforms SOTA methods on four prediction tasks. **(A)**. ROC curves of ESM-DBP and SOTA methods on four downstream prediction tasks; **(B)**. MCC values of ESM-DBP and SOTA methods on three downstream prediction tasks; **(C)**. Head-to-head comparisons of MCC values between ESM-DBP and other SOTA methods on PDNA-41. Each point means a DBP and the axes indicate the MCC value of the predictor on this protein. The color bar indicates the amount of DBS on the DBP; **(D)**. The prediction results of ESM-DBP and SOTA methods on the chain C of a trimer (PDB ID: 4ZCF) with an MSA depth of 253 against Uniclust30 database [41] using HHblits program [35].

Figure 3A shows the ROC curves of ESM-DBP and the SOTA methods on four prediction tasks. What is clearly visible is that the curve of ESM-DBP is significantly higher than the other predictors, regardless of the task. For instance, the AUC value of ESM-DBP for DBS prediction is 0.905, which achieves improvements of 6.59, 8.77, 21.47, and 9.30% over iDRNA-ITF, NCBRPred, DRNAPred, and CLAPE-DB separately. Considering the MCC metrics (shown in Figure 3B), ESM-DBP (highlighted with lavender) also achieves the best results over three tasks. Notably, the MCC of ESM-DBP on TF prediction exceeds 0.9, which is much higher than the other three methods (none of them exceeded 0.75). The main reason is that the specific sequences of TF possess a high degree of conservatism [1] easily detected by PLM to give more attention. Figure 3C illustrates the head-to-head comparison of MCC values among ESM-DBP and other methods over the DBS predictions. In most cases, ESM-DBP archives a higher MCC value compared to either method. Specifically, out of 41 DBPs, there are 31, 32, 36, and 25 cases in which ESM-DBP outperforms iDRNA-ITF, NCBRPred, DRNAPred, and CLAPE-DB, respectively. Of these four methods, the first three are evolutionary information-based methods, and the latter, like ESM-DBP, is a PLM-based method and fine-tuned based ProtTrans [23]. Here, the chain C of a trimer (PDB ID: 4ZCF) with an MSA depth of 253 against Uniclust30 is employed as a case study to demonstrate the performance of these methods on sequences with sufficient MSA information. As shown in Figure 3D, it is clear that the three methods based on evolutionary information do not make a good prediction. Potential reasons for this are the complexity of the structure of trimers and DNA complexes, as well as long amino acid sequences that limit the predictive accuracy of these methods. In contrast, as the PLM-based prediction methods, CLAPE-DB (MCC: 0.159) and ESM-DBP (MCC: 0.440) performs better than three evolutionary information-based methods, which demonstrates the DBS prediction ability of PLM for large complexes. In this case, the excellent performance of ESM-DBP compared to CLAPE-DB may come from retraining on the DBP sequence data, making ESM-DBP more sensitive to DNA-binding regions. In summary, ESM-DBP demonstrates excellent prediction performance over all four prediction tasks and outperforms existing methods at the SOTA level.

### ESM-DBP explores the DNA-binding domain

The ability of a protein to bind to DNA mainly comes from the DNA-binding domain (DBD) such as HMG box, Homeobox, Fork-head, and Nuclear receptor. Even for transcription factors, the ability to bind specific DNA sequences alone is usually a reflection of the ability to regulate transcription [1]. ESM-1b reports on the capability of PLM to explore the remote homology and the MSA information in the protein family of protein sequence. The pretraining dataset containing 170,264 DBP in this study has an abundance of DBD sequences. In general, those protein sequences that contain DBDs of the same class generally have higher homology and closer evolutionary distance. Theoretically, PLM can distinguish between DBPs and non-DBPs by capturing the commonality of DBDs of these similar DBP sequences. Here, to validate this idea, we first feed 142,657 single-domain protein sequences in UniDBP40 whose domain data can be found in UniProt into the ESM-DBP model to derive sequence embedding features; then, T-SNE algorithm is used to visualize these embedded features. From Figure 4A, for those families with abundant sequences like HTH and Homeobox, clustering is effective and can be clearly distinguished. Domain-adaptive pretraining allows PLM to pay more attention to those families of proteins with high frequency of occurrence than to common proteins. In addition, for the TFs in the test set, we use the integrated gradient algorithm [58] to observe the contribution value of the residues at each position in the sequence to the prediction result (see https://github.com/pengsl-lab/ESM-DBP). In Figure 4B, five single DBD human TFs (UniProtKB ID: P35713, Q9UBX0, P58021, Q9NX45, and Q10587) separately containing five typical DBDs, i.e., HMG box, Homeobox, Fork-head, bHLH, and bZIP, as well as one multi-DBD TF (UniProt ID: P51449) containing two domains, i.e., Nuclear receptor and NR LBD, are used as demonstration cases. By looking at Figure 4A, the most intuitive feeling is that those residues in the DBD region have significantly higher saliency scores than those in the non-DBD region regardless of protein. For instance, residues between index 85 and 153 on protein P35713 are significantly darker in color than the rest of the residues and look linear. This suggests that residues in the DBD region contribute particularly to the prediction results, and it is because of the perception of this sequence fragment that PLM identifies TF. We also note that not all residues in the DBD region have significant saliency scores; in fact, only a small number of residues have a large effect, and in particular, only one residue is more prominently colored on Q10587. Moreover, the contribution of a residue in a non-DBD region is also evident on Q9UBX0. After all, DBD and non-DBD of a protein sequence do not exist in isolation but have an interactive relationship. Residues in the non-DBD region also may affect the binding pattern to DNA, although this effect is much less than that of residues in the DBD region in most cases. From the results of the multi-domain protein P51449, residues within both domains positively influence the final prediction, indicating that ESM-DBP not only probes the knowledge of single-domain proteins but also captures the specificity information of different DBDs of multi-domain proteins.

**Figure 4.**
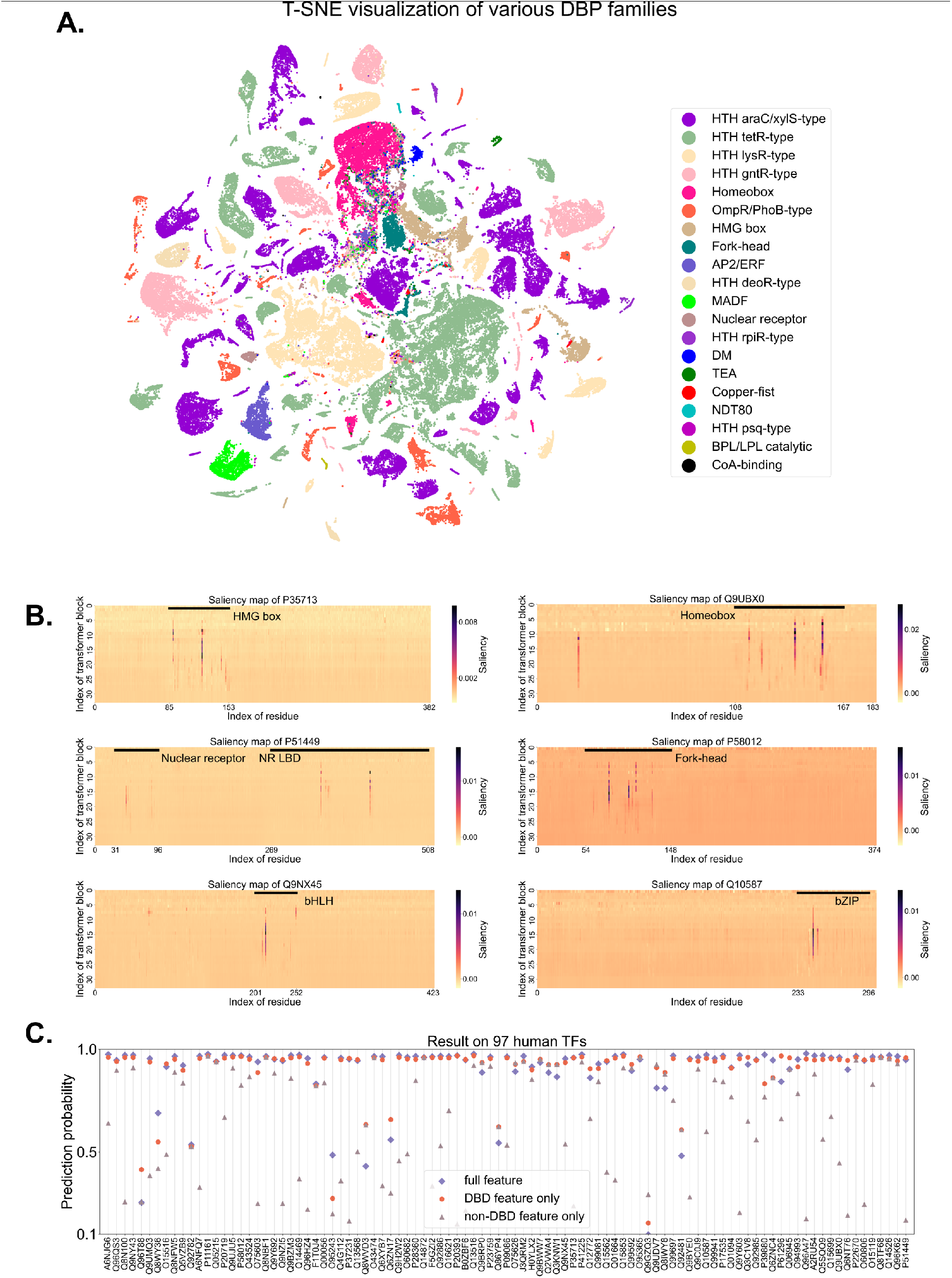
ESM-DBP explores the DNA-binding domains. **(A)**. Visualization of ESM-DBP sequence embedding of different DBD families in UniDBP40; **(B)**. The saliency map of six human TFs. The solid black line indicates the DBD area recorded in UniProt. The horizontal axis is the index of the residue in the protein sequence. The vertical axis represents the index of the transformer block in ESM-DBP. Each value with coordinates (*x, y*) represents the saliency score calculated by the integrated gradient algorithm of the *x*-th residue in the *y*-th transformer block for the prediction result; The integrated gradient algorithm is implemented in the Captum library [57]; **(C)**. Prediction results comparison of full feature (marked in diamond), DBD feature (marked in the circle), and non-DBD feature (marked in the triangle) on 91 test human TFs. The DBD (or non-DBD) feature is obtained by replacing all residues that are not in (or in) the DBD region with the token of <mask> before inputting protein sequence into the ESM-DBP model.

To further demonstrate the role of DBD on the predictive performance of ESM-DBP, we compare the performance of three different features on 97 test human TFs, which are obtained by inputting the complete protein sequence, the sequence of the DBD region, and the sequence of the non-DBD region into ESM-DBP, respectively. In Figure 4C, we can clearly observe that in most of the cases, the prediction probability of the non-DBD features drops drastically compared to the full features since the critical DNA binding pattern identification information is missing from the feature after masking out the sequences in the DBD region. In contrast, the prediction results using DBD features are not nearly as different from the full features since the integrity of the identification information in the DBD region. Undoubtedly, the inference of ESM-DBP depends greatly on the sequence fragments of the DBD region. To sum up, ESM-DBP perceives various DBDs hiding DNA binding patterns from a large number of DBP sequences in the pre-training dataset, which enables accurate recognition of DBPs and TFs.

### ESM-DBP is good at generalizability

In the section “ESM-DBP explores the DNA-binding domains”, we mention that ESM-DBP learns specific DBD knowledge from clusters of similar protein sequences to improve the prediction performance of related tasks. In nature, some orphan proteins with a short evolutionary history lack homologous proteins. This is the main reason that limits the performance of most protein function prediction methods including AlphaFold2. Here, we observe the relationship between sequence similarity of test protein against to pretraining data set and prediction performance. Specifically, for the 381 positive samples in the DBP test set, UniDBP40 and UniRef50 are used as target databases to search for homologous sequences using the blast program with an e-value of 1e-06, respectively (results see https://github.com/pengsl-lab/ESM-DBP). By visiting Figure 5A, out of 381 DBPs, most of the proteins have only a small number of high similarity sequences against UniDBP40, moreover 155 cases have no high similarity sequences; due to the abundance and diversity of sequences in UniRef50, a high number of similar sequences can be found for most of the test proteins. Eighteen DBPs (marked in the pentagram) with a prediction difference between ESM-DBP and ESM2 greater than 0.4 are present in the gray highlighted region of Figure 5B, most of which have few similar sequences with UniDBP40. In Figure 5C, looking at the proteins that have abundant similar sequences against UniDBP40 and UniRef50 respectively (located in the upper and right halves respectively), all proteins in the former have good predictions, while some proteins with poor predictions exist in the latter. Considering the excessive number of sequences in UniRef50, ESM2 could not adequately focus on all DBPs. In contrast, ESM-DBP mined as much valid information as possible from UniDBP40 by domain-adaptive pretraining. Looking closely at those proteins at the bottom with less sequence similarity to UniDBP40, most of them still have good predicted probability values. Figure 5D shows the predicted results of 155 DBPs without any similar sequences and 226 DBPs in the presence of homology proteins against UniDBP40 respectively. From the two sets of box plots at the top of Figure 5D, both ESM2 and ESM-DBP have much higher predicted probabilities on high homology proteins than those on low homology proteins, since the model detects those similar sequence fragments many times during the pretraining period and thus gives the more attention to them. Comparing the results of ESM2 and ESM-DBP on low homology proteins, the overall distribution of ESM-DBP is significantly higher than that of ESM2. These phenomena demonstrate that ESM-DBP mines key knowledge from those DBPs with only a small number of similar sequences, allowing it to further enhance the DNA-binding pattern characterization based on ESM2 for better prediction.

**Figure 5.**
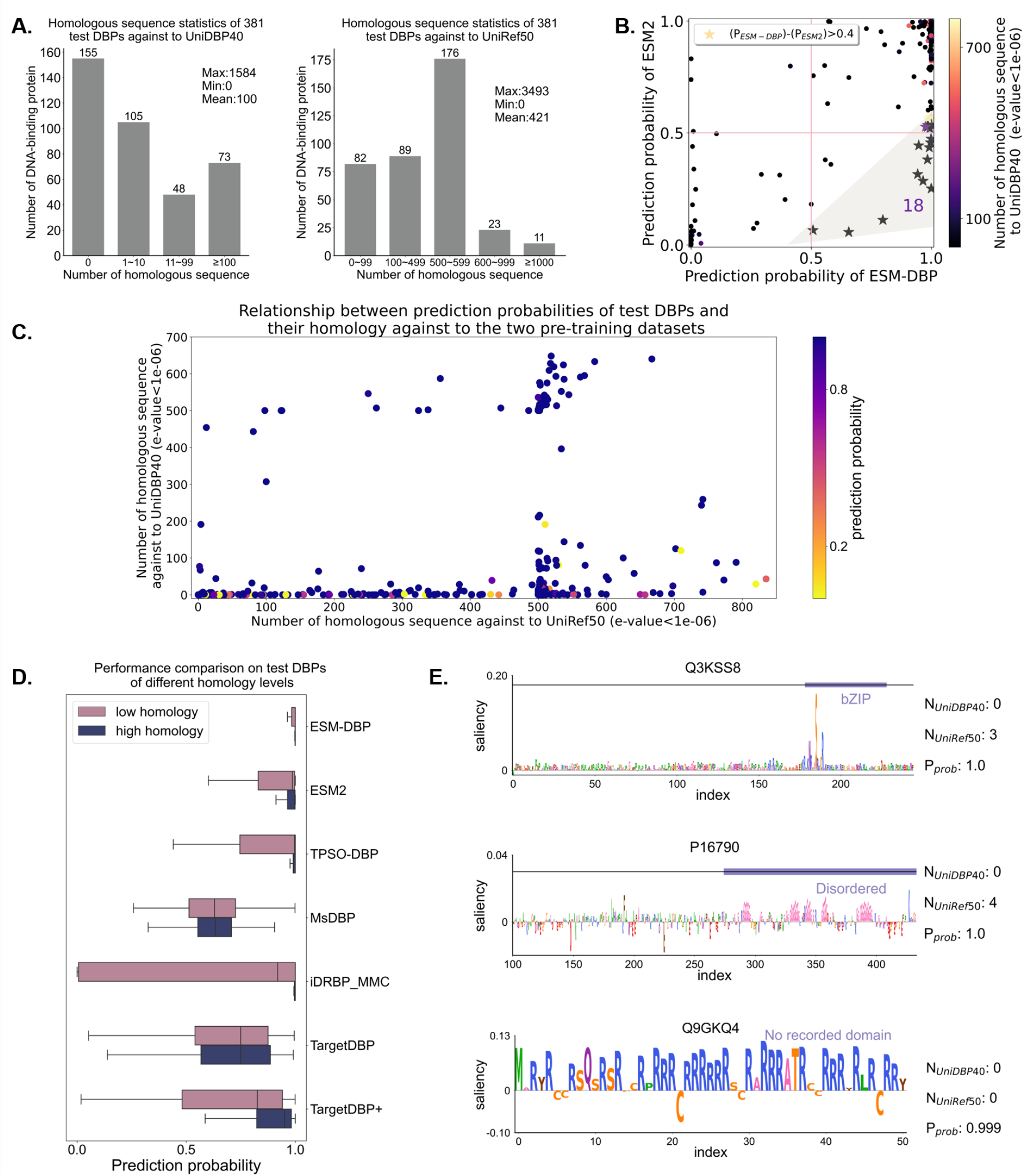
ESM-DBP performs well on DBPs with few homologous proteins. **(A)**. Statistics on the number of homologous proteins of 381 DBPs in the independent test set against UniDBP40 and UniRef50 respectively using the blast program with an e-value of 1e-06; **(B)**. Head-to-head comparison between the prediction probabilities of ESM-DBP and ESM2 on the 381 DBPs in the independent test set. Pentagrams highlighted in grey shading indicate those 18 DBPs for which the difference between the predicted probabilities of ESM-DBP and ESM2 is greater than 0.4; **(C)**. The relationship between the prediction probability of DBP and its homology against the two pre-training datasets; **(D)**. Prediction results of ESM2 and other methods on 226 high (or 155 low) homology DBPs in independent test set whose number of similar sequences against UniDBP40 greater than 0 (or equal to 0); **(E)**. ESM-DBP learns discriminatory knowledge from three different types of low homologous proteins, i.e., bZIP protein, disordered protein, and protein without domain. N_*UniDBP40*_ and N_*UniRef50*_ mean the number of similar sequences against to UniDBP40 and UniRef50 separately. P_*prob*_ indicates the probability of being predicted as a DBP by ESM-DBP.

We also compared with the SOTA DBP prediction methods that mostly rely on high quality MSA information on test DBPs with different homology levels. (see Figure 5D). Obviously, most other predictors show a significant decrease in prediction scores on low homology DBPs compared to high homology DBPs. The limited available evolutionary information is the main reason that constrains the generalizability of these traditional methods. In contrast, after continued training and full exploration on a large number of DBP sequences, ESM-DBP is able to capture valid discriminatory information about DBPs that have even only a small number of homologous sequences. The outstanding performance of ESM-DBP on these DBPs further demonstrates the generalizability that other methods lack. In Figure 5E, three cases (UniProt ID: Q3KSS8, P16790, and Q9GKQ4) are employed to demonstrate the ability of ESM-DBP to learn the discriminatory information of various low homology proteins. The saliency of the amino acid residue at each position indicates the contribution of that residue to the final prediction. The first two cases contain the bZIP domain and the disorder region, respectively; the latter has no domain recorded in the UniProt database. The saliency maps for the first two cases show the high sensitivity of ESM-DBP to bZIP domain and disorder region. The bZIP is a typical DBD common among transcription factors in eukaryotes. Previous studies have reported the important effects of intrinsically disordered regions on DNA-protein interactions [59-63] and the C-terminal region of P16790 (DNA polymerase processivity factor) is required for viral DNA synthesis [64]. The specific recognition of both highlights the capability of ESM-DBP to draw valid knowledge from protein domain in low homology sequences. The latter exemplifies the ability of ESM-DBP to identify DBP even in the absence of typical DBDs in the orphan protein. The positive saliency score for the vast majority of residues in the entire sequence highlights the ability of ESM-DBP to capture recognition knowledge from global sequence rather than a local, specific protein domain. These results firmly support the idea that ESM-DBP can learn DBP discriminative information for proteins with few homologous sequences. The generalization ability demonstrated by ESM-DBP provides a potential new perspective for solving the problem of structure and function prediction of orphan proteins.

## CONCLUSIONS

Protein language models have demonstrated superior learning capabilities for the representation of biological knowledge, leading to great advances in protein structure and function prediction. However, due to the extremely complex and nonlinear relationship between protein sequence, structure, and function, the characterization of sequences determining a particular biological function by the general protein language model has yet to be improved. In this study, we propose ESM-DBP, a DNA-binding protein field-specific protein language model based on ESM2, which is constructed by domain-adaptive pretraining on massive non-redundant DNA-binding protein sequences. The prediction results on the four DBP-related downstream tasks show that ESM-DBP achieves a better characterization of DBP sequences than ESM2, which improves the prediction performance. Meanwhile, the prediction accuracy of ESM-DBP on all four tasks far exceeds that of the state-of-the-art prediction methods.

In the future study, we will focus on the following points: (1). extending the idea of domain-adaptive pretraining to other protein function studies like RNA-binding protein and protein-protein interaction; (2). constructing multi-modal large biologic language model from proteins and other ligands such as ATP and RNA to further improve the sequence feature representation; (3). achieving the end-to-end protein-nucleic acid complex structure prediction using a multi-modal language model.

This study suggests that the sequence characterization capability of a generic large-scale protein language model can be further explored, providing a new perspective on the protein function study. In addition, ESM-DBP shows outstanding prediction performance for DBP-related tasks, thus accelerating the process of protein-DNA interaction research.

## DATA AVAILABILITY

The data, source code, and standalone program of ESM-DBP are freely accessible at https://github.com/pengsl-lab/ESM-DBP. The pretraining data set and ESM-DBP models are available for academic use at https://huggingface.co/zengwenwu/ESM-DBP/tree/main.

## AUTHOR CONTRIBUTIONS

Wenwu Zeng: Conceptualization, Methodology, Software, Visualization, Writing original draft. Yutao Dou: Methodology, Software, Visualization. Liangrui Pan: Conceptualization, Visualization. Liwen Xu: Conceptualization, Methodology, Visualization, Writing original draft. Shaoliang Peng: Funding acquisition, Resources, Writing-review × editing, Supervision.

## ACKNOWLEDGMENT

We would like to thank the Funds of State Key Laboratory of Chemo/Biosensing and Chemometrics, the National Supercomputing Center in Changsha (http://nscc.hnu.edu.cn/), and Peng Cheng Lab. L.W. Xu, and S.L. Peng are the corresponding authors for this paper.

